# A nucleation distance sets the cell size-dependent P-body assembly

**DOI:** 10.1101/2025.07.08.663506

**Authors:** Xuefang Gu, Dexin Shen, Xing Quan, Tatsuhisa Tsuboi

## Abstract

Cells rely on diverse intracellular structures, including phase-separated granules, to regulate biological processes. Processing body (P-body) is a type of membraneless condensate in the eukaryotic cytoplasm that mediates mRNA degradation and storage. While their molecular composition is well studied, the relationship between their spatial organization and cell geometry, particularly cell size and organelle distribution, remains unclear. Here, we develop a conceptual framework that models phase-separation nucleation and validates the nucleation distance of P-bodies in living yeast cells. Using microfluidic cell sorting and quantitative fluorescence microscopy, we find that small cells contain fewer but larger P-bodies, whereas large cells form more but smaller granules. Monte Carlo simulations of P-body nucleation accurately recapitulate these cell size-dependent patterns and reveal a nucleation distance of 2.2 µm. Furthermore, we demonstrate that increasing vacuolar volume reduces the effective cytoplasmic space and correspondingly raises P-body number, although cell size remains the dominant determinant. Our findings establish nucleation distance as a key parameter that links cell geometry to the biophysical regulation of condensate formation.

## Introduction

Cellular structures act in concert to regulate essential processes such as intracellular transport, energy conversion, and gene expression ^1–3^. Among these structures, membrane-less organelles formed through liquid-liquid phase separation (LLPS) play a critical role in gene expression regulation ^4–6^. Cytoplasmic Processing bodies (P-bodies) are prototypical LLPS condensates in eukaryotic cells ^3^. They sequester translationally repressed messenger RNAs (mRNAs) together with the decapping and decay enzymes ^7,8^, thereby orchestrating mRNA degradation, storage, and recycling ^8–11^. P-body assembly involves several partially redundant scaffold proteins, including Edc3^12,13^, Dcp2^14^, Dhh1 ^15^, Pat1 ^15,16^, and Lsm4 ^12,17^. Deletion of these key proteins typically reduces P-body number and size and impairs the recruitment of client RNAs and proteins ^18,19^.

Phase separation begins with nucleation, the process in which a small molecular cluster becomes large enough to overcome the energetic cost of forming a new interface. In supersaturated solutions or vapours, only molecules that diffuse within a reaction radius can bind productively; beyond this distance, collisions are too infrequent to surpass the nucleation barrier ^20^. Once a condensate forms, it recruits nearby molecules, creating a depletion zone that reduces local supersaturation. New droplets can only nucleate beyond this depleted region. Together with the natural tendency of neighboring droplets to fuse and minimize surface energy, this phenomenon defines a characteristic nucleation distance (dₙ), the typical spacing at which independent condensates can stably coexist ^21^. Inside cells, molecular crowding and sub-diffusive dynamics further limit droplet fusion, allowing many condensates to persist at approximately dₙ apart rather than merging into a single large phase ^22^. Simulations using a coarse-grained protein model reveal that the stability of biomolecular condensates depends mainly on the number of attractive interactions between molecules ^23^. By modulating concentration, crowding, or binding affinity, cells can tune dₙ to control both the number and spatial distribution of condensates.

Despite the complexity of the intracellular environment, nucleation in living cells still adheres to fundamental physics constraints, the existence of an energy barrier, and the requirement for a critical molecular cluster, with cellular components often acting as seeds to facilitate this process^20^. The concept of a nucleation distance thus bridges physical theory and cell biology, as it arises from the necessity of accumulating sufficient molecules within a reaction radius and is reinforced by molecular depletion effects and coalescence dynamics, ultimately leading to a regulated, non- random spatial distribution of condensates within the cell. These insights suggest that P-body assembly is governed not only by protein composition and interactions but also by the physical dimensions and spatial organization of the cell.

Cellular organization is determined by the geometry of intracellular compartments and directly influences the scale and efficiency of biosynthetic processes ^24–26^. As cells grow larger, organelles generally scale proportionally ^27–29^. Although the total protein concentration is maintained, the concentration of individual proteins may increase or decrease with cell size ^30–32^. Interestingly, P- bodies are known to appear during G1 and expand in size through G2, without a corresponding change in number ^33^, suggesting that spatial constraints, rather than protein abundance, may play a key role in phase separation. Yet, the precise influence of cell size and spatial parameters on P- body formation and dynamics remains largely unexplored.

Here, we extend this conceptual framework to quantitatively model condensate nucleation and validate the presence of a characteristic nucleation distance in living cells. By combining quantitative microscopy of size-sorted cells using microfluidic devices with Monte Carlo simulations for nucleation, we find that larger cells tend to form more but smaller P-bodies, whereas smaller cells generate fewer but larger P-bodies. The simulations identify a specific nucleation distance (dₙ) that aligns closely with our experimental observations. Moreover, in *sch9*Δ and *vps1*Δ mutants, where vacuolar volume is enlarged and the effective cytoplasmic space reduced, we observe a significant increase in P-body number, reinforcing the importance of spatial restriction in phase separation. Together, these results enable us to investigate how cells regulate the nucleation distance to influence the formation, size, and distribution of phase-separated compartments.

## Results

### P-body number increases with yeast cell size

To examine the relationship between cell size and P-body formation under physiological conditions, we quantify how the spatial organization and cell geometry of a yeast cell influence the initiation and abundance of P-bodies *in vivo* while avoiding genetic manipulations that can inadvertently perturb LLPS ^34,35^. We developed a microfluidic device based on cell trapping for long-term time-lapse microscopy ^36,37^, containing one hundred parallel trap lines whose constriction widths increase stepwise from 5.1 µm to 10 µm, enabling continuous perfusion of logarithmically growing W303 cells and deterministic physical sorting by diameter (Fig. 1A, B, Supplementary Fig. 1). This non-invasive, high-throughput platform let us examine a wide range of cell sizes in a single experiment while minimizing batch-to-batch variability (Supplementary Fig. 1C). This platform preserved nutrient availability while imaging under an identical glucose-starvation regime (0 % glucose, synthetic complete medium), a known physiological stress that robustly induces P-body formation. The use of physical sorting rather than genetic perturbation was essential to isolate the impact of cell size and to avoid confounding factors such as stress responses, altered expression of phase-separation components, or cell-cycle defects.

**Figure 1.**
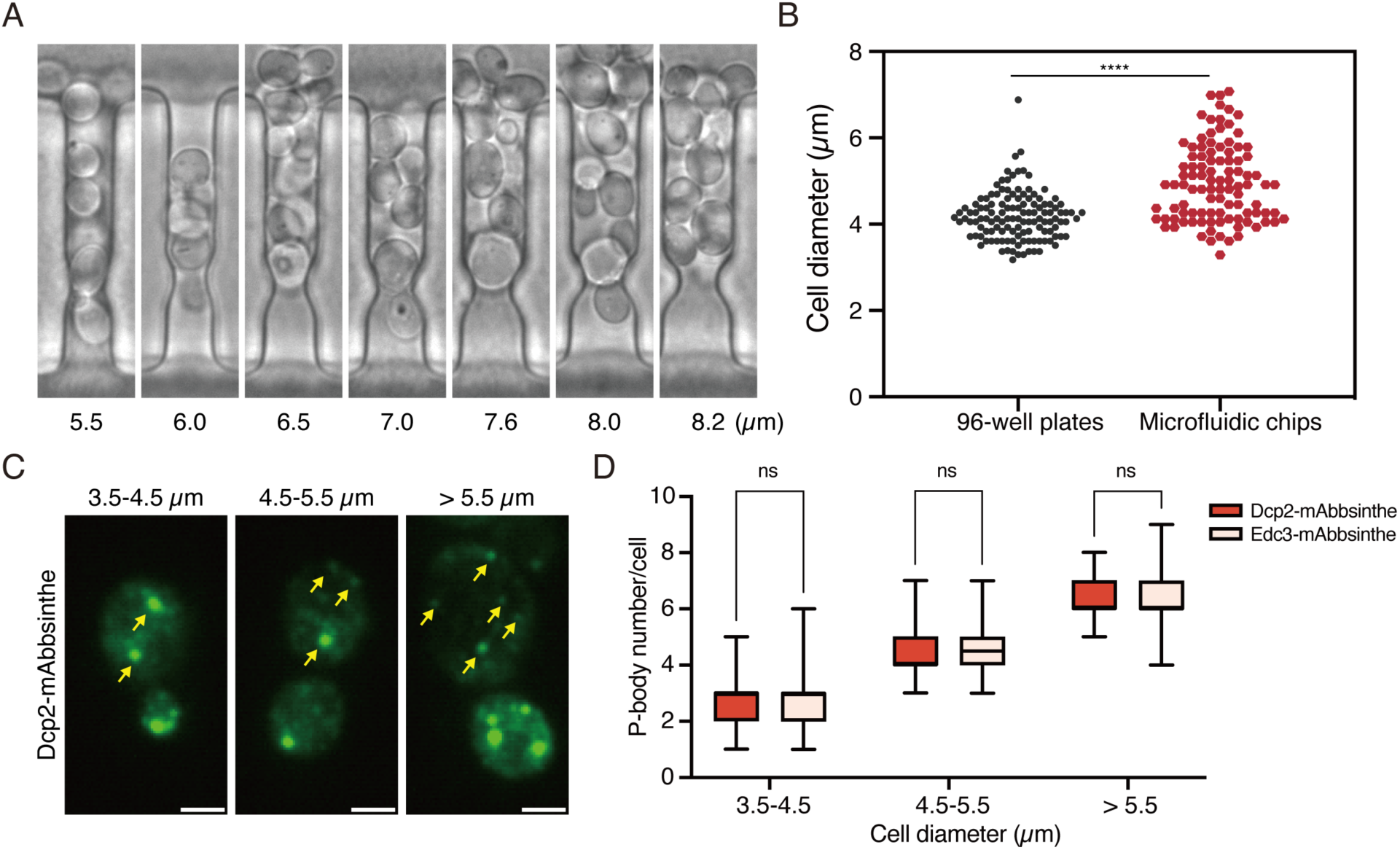
| P-body number increases with increasing yeast cell size (A) Yeast cells of different sizes are captured in the microfluidic device; trap widths are indicated below each channel. (B) Comparison of cell diameters in 96-well plates and microfluidic chips. Each dot corresponds to a single cell (n>90). Statistical significance was assessed by the Unpaired T-test (**** P < 0.0001). (C) P-bodies in varying sizes of yeast cells were labeled with Dcp2-mAbbsinthe and analyzed 30 minutes after glucose starvation. Yellow arrows indicate P-bodies in mother cells. Scale bar, 2 µm. (D) P-body number per cell across different cell sizes. Yeast cells expressing Dcp2-mAbbsinthe or Edc3-mAbbsinthe were categorized into three size ranges based on cell diameter. Statistical significance was assessed by Unpaired T-test (ns P > 0.05, n>30).

We tagged Dcp2, a core P-body scaffold, with a newly engineered green fluorescent protein variant, mAbbsinthe, which exhibits superior photostability and lower self-aggregation relative to conventional GFPs (Supplementary Fig. 2A, B)^38^. Dcp2-mAbbsinthe retained its ability to induce P-body formation in response to stress, confirming its suitability as a functional reporter (Supplementary Fig. 2C). The enhanced photostability of mAbbsinthe was critical for long-term imaging of condensate dynamics under glucose starvation. Using spinning-disk confocal microscopy, we acquired three-dimensional stacks of P-body distribution in yeast cells spanning the full diameter range (Fig. 1C). Quantitative analysis revealed a strong positive correlation between cell size and p-body number (Fig. 1D). The mean P-body number per cell rose from 2.7 in 3.5–4.5 µm cells to 4.4 in 4.5-5.5 µm cells, and to 6.2 in cells larger than 5.5 µm. These findings were reproduced with Edc3-mAbbsinthe, an alternative P-body marker, demonstrating that the effect is not reporter-specific. Together, our results provide direct experimental evidence that cell size is a key determinant of P-body number, suggesting that cellular geometry influences phase separation behavior *in vivo*.

### A Monte-Carlo simulation based on nucleation identifies a characteristic nucleation distance

To test whether simple geometric constraints could explain the empirically observed scaling, we implemented a Monte-Carlo simulation based on nucleation. In this model, we approximated the yeast mother cell as a sphere and randomly distributed virtual Dcp2 molecules within its volume (Fig. 2A). This abstraction was motivated by the physical properties of phase-separated condensates, which behave as mesoscale entities with restricted mobility and volume-excluding effects. Collisions resulting in center-to-center separations below a tunable threshold (nucleation distance, dₙ) trigger irreversible coalescence, conserving total mass. These fusion events were iteratively calculated until all remaining molecular pairs were spaced beyond this threshold. We initialized the model with 200 seed spheres per cell, each with a radius of 2.0 to 3.0 µm. Across this range, a best-fit dₙ of 2.2 µm simultaneously reproduced the monotonic increase in puncta number with cell size (Fig. 1D, 2B, Supplementary Figure 3A). We then compared the size of P- bodies between simulations and experiments. Since P-bodies were visualized through microscopy imaging, apparent measurements can deviate from actual biological sizes. To ensure consistent comparison, we quantified P-body size by calculating the proportion of puncta in each size category (Supplementary Figure 3B). We first matched the category proportion between the two experiments for a cell size of 5.0 µm. With this size classification, the simulation for the other two different size cells, 4.0 µm and 6.0 µm, matches surprisingly well with the empirically observed triphasic size distribution of P-bodies (Fig. 2C - G). Smaller cells tend to form larger P-bodies, whereas larger cells predominantly contain smaller P-bodies. This agreement suggests that the physical spacing between successful nucleation events, a feature not encoded in protein sequence or interaction affinity, acts as a central organizing principle. To probe robustness, we performed sensitivity analyses that varied initial molecular numbers and cytoplasmic viscosity (Supplementary Figure 3C, D). The final puncta number was largely invariant to particle count above ∼200 and only weakly dependent on viscosity (within biologically plausible ranges), indicating that dₙ is the dominant determinant of condensate distribution. These simulations highlight that spatial constraints and collision thresholds alone are sufficient to explain the observed morphological transitions, without invoking complex biochemical regulation.

**Figure 2.**
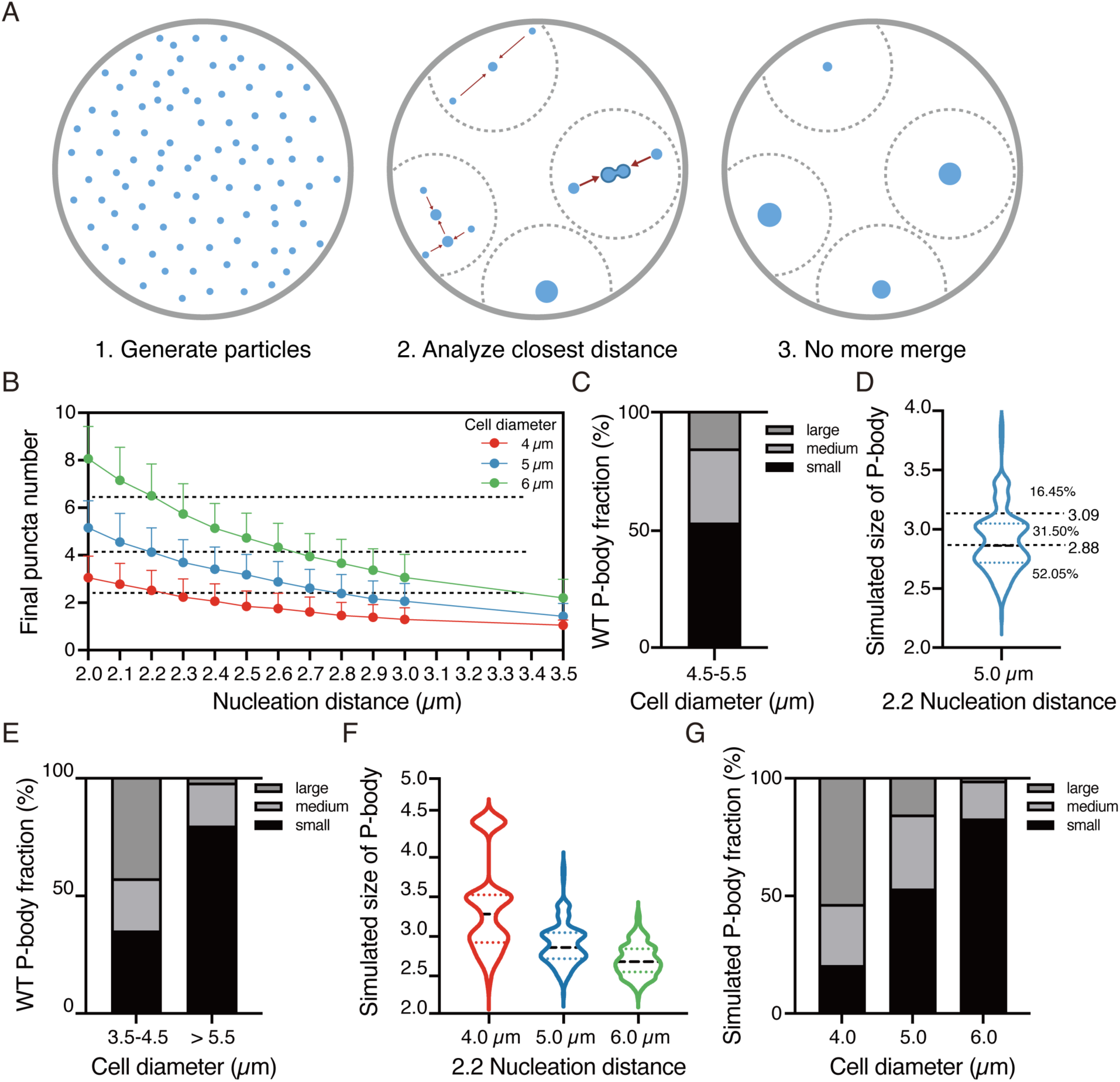
| A Monte-Carlo simulation based on nucleation identifies a characteristic nucleation distance (A) Schematic representation of particle nucleation model based on Monte Carlo simulation. 1. Generate particles: Initial distribution of molecules in 3D space before simulation, each blue dot represents a molecule. 2. Analyze the closest distance: Each molecule recognizes the closest one and then fuses. 3. No more merge: Result of the nucleation model, showing puncta left after merge, each granule represents a P-body. (B) Nucleation distance correlates with the final puncta number. Based on the nucleation distance, final puncta numbers were simulated for each cell size (n = 200 per condition). Horizontal dashed lines indicate the mean P-body number per cell of different cell size groups in Figure 1D. (C) Size fraction of P-body in the 4.5-5.5µm cell size by the manual classification of the microscope image. (D) The definition of the size of the simulated P-bodies. The boundary for different sizes of P- bodies in the simulation is based on the fraction in C. (E) Size fraction of P-body in the different cell sizes by the manual classification of microscope image data. (F) Size of P-bodies in different cell sizes at the dₙ = 2.2 µm in the simulation, including D. (G) Fractions of the simulated P-body size in the different cell sizes based on the definition in D.

### Organelle occupancy amplifies nucleation frequency by restricting effective cytoplasmic volume

While total cell volume increases during growth, the effective cytoplasmic space available for P-body formation may be limited by the presence of large organelles such as the vacuole and nucleus. To explore the impact of such intracellular crowding, we extended our simulation to incorporate internal organelles occupying specific volumetric fractions. We implemented three distinct models to simulate molecule fusion paths in the presence of a central spherical nucleus and vacuole, representing non-permissive organelle space: (1) path-restricted fusion, in which straight-line paths that crossed the nucleus were invalid, (2) path-shifted fusion, in which granules repositioned to avoid organelle overlap, and (3) path-redirected fusion, in which particles navigated around obstacles using a calculated shortest path (Fig. 3A–C).

**Figure 3.**
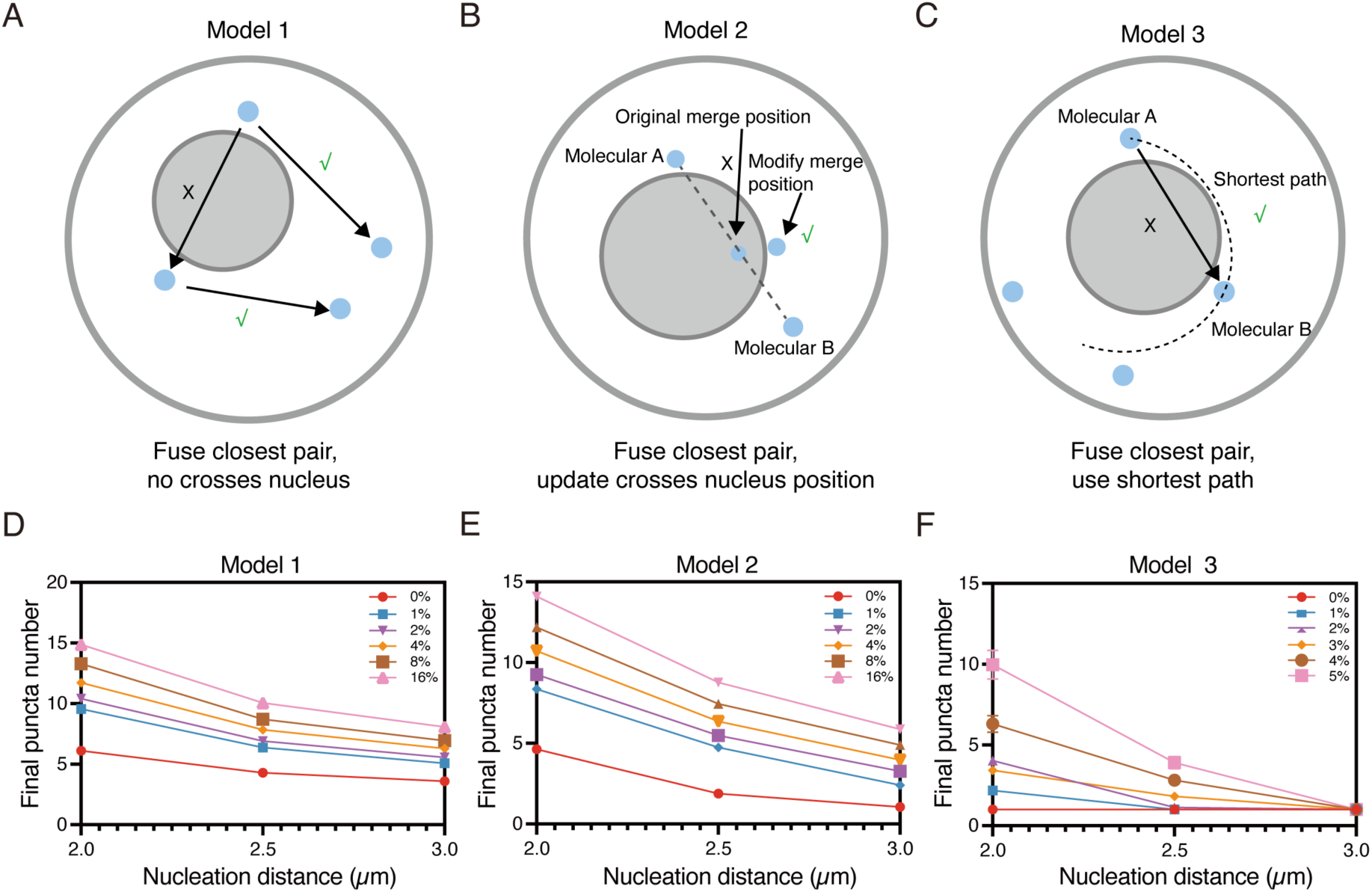
| Organelle occupancy amplifies nucleation frequency by restricting effective cytoplasmic volume (A-C) Schematic diagrams of three nucleation models in a cell containing a centrally located organelle (gray circle). The symbols “X” and “√” show whether a selected fusion path is valid based on each model’s rules. “X” indicates it is not allowed, while “√” signifies it is permitted. (A) Model 1: Molecules fuse with their closest neighbor only if the connecting line does not intersect the central organelle. (B) Model 2: Molecules fuse with their closest neighbor even if the direct path crosses the nucleus, but the merged granule is repositioned to avoid overlap with the central organelle. (C) Model 3: Molecules fuse with their closest neighbor based on the true shortest path around the nucleus, allowing nucleus-aware trajectories. (D-F) Final puncta number as a function of nucleation distance and organelle volume fraction. Color represents the different organelle fractions relative to the cell.

Across all models, increasing organelle volume fractions consistently led to elevated final P-body counts, even under a constant nucleation distance (Fig. 3D–G). This effect arose because internal obstructions constrained the diffusion space, increasing local molecule concentrations and reducing the effective reaction volume. These simulations suggest that intracellular crowding can enhance condensate nucleation independent of changes in protein concentration or binding affinity, highlighting a potential mechanism for tuning granule formation through organelle scaling or spatial reorganization.

### Vacuole mutants validate the influence of spatial confinement on P-body assembly

To test the simulation-derived hypothesis that intracellular crowding promotes condensate nucleation, we examined two yeast mutants known to affect vacuolar morphology: *sch9*Δ and *vps1*Δ. Both mutations increase vacuole-to-cell volume ratio through distinct mechanisms: *vps1*Δ via impaired membrane fission and *sch9*Δ via defects in nutrient sensing and vacuolar growth regulation ^27,39^. Quantification of vacuole-to-cell volume ratios confirmed significantly increased organelle occupancy in both mutants compared to wild-type (Fig. 4A, B). Fluorescence intensity analysis showed that Dcp2-mabbsinthe expression remained unchanged (Fig. 4C), ensuring that any observed effects on P-bodies were not due to altered protein abundance. As predicted by the simulation, *vps1*Δ cells exhibited significantly higher P-body counts across all size bins (Fig. 4D), indicating that reduced cytoplasmic space can elevate nucleation rates. Surprisingly, *sch9*Δ cells, despite also showing increased vacuole volume, did not exhibit a similar rise in P-body number. Instead, both *sch9*Δ and *vps1*Δ displayed a marked shift toward smaller P-body sizes (Fig. 4E, F), especially in larger cells.

**Figure 4.**
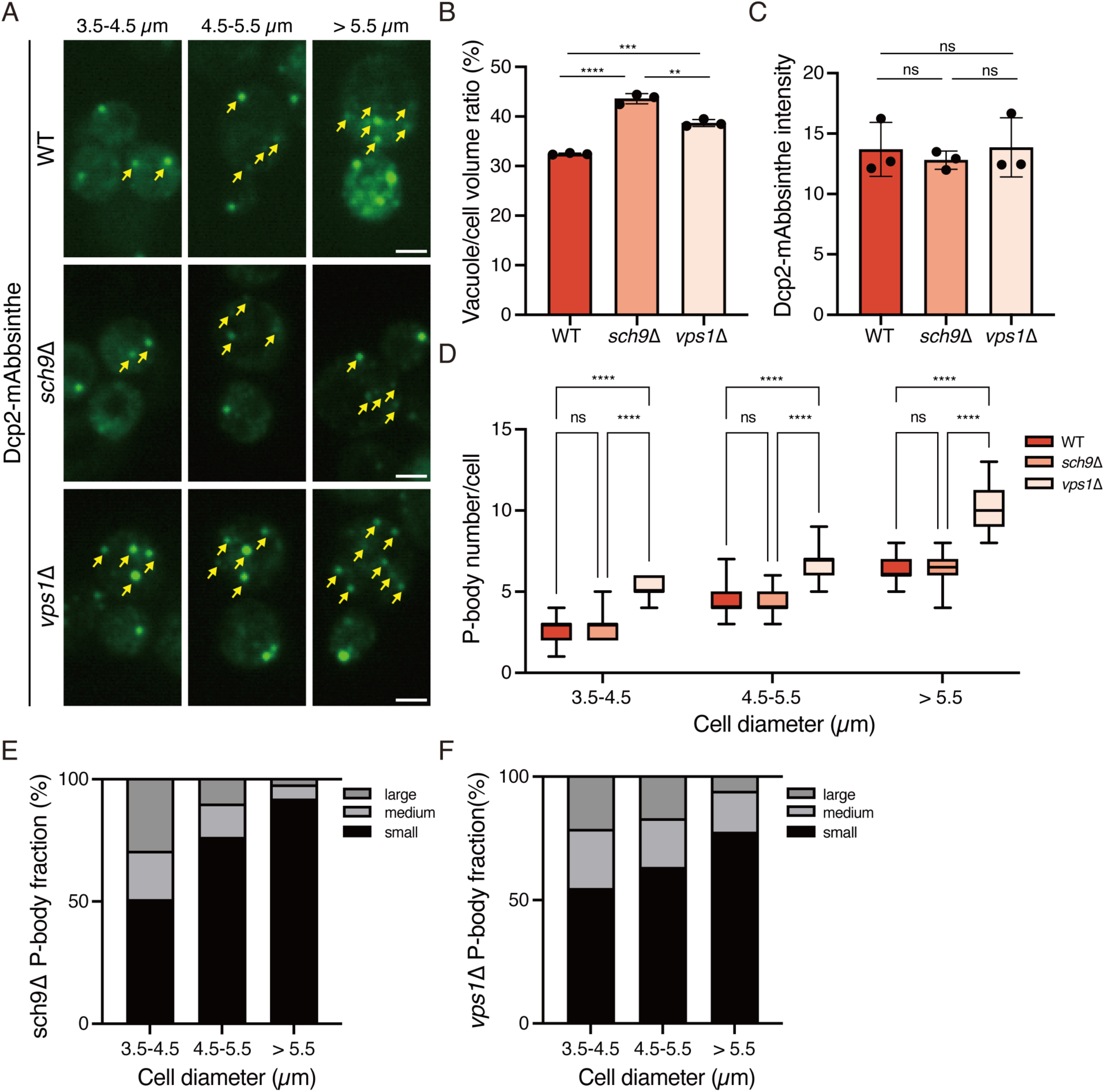
| Vacuole mutants validate the influence of spatial confinement on P-body assembly (A) Visualization of P-bodies in WT, *sch9*Δ, and *vps1*Δ yeast strains expressing Dcp2-mAbbsinthe under glucose starvation for 30 minutes. Cells are grouped by cell diameter (3.5-4.5 µm, 4.5-5.5 µm, >5.5 µm), and P-bodies are indicated by arrows. Images are z-projections of 42 slices. Scale bar, 2 µm. (B) Quantification of vacuole-to-cell volume ratio in WT, *sch9*Δ, and *vps1*Δ strains. Each dot represents an individual experiment (n = 3 biological replicates). Statistical analysis was performed using one-way ANOVA with multiple comparisons (**P < 0.01, ***P < 0.001, ****P < 0.0001). (C) Mean Dcp2-mAbbsinthe fluorescence intensity in WT, *sch9*Δ, and *vps1*Δ strains. No significant differences were observed. Each dot represents an individual experiment. (D) P-body number per cell across different cell diameter groups (3.5-4.5 µm, 4.5-5.5 µm, >5.5 µm) in WT, *sch9*Δ, and *vps1*Δ strains. Statistical significance was determined by two-way ANOVA with multiple comparisons (ns P > 0.05, ****P < 0.0001). (E) Proportion of large, medium, and small P-bodies in *sch9*Δ cells across the cell size groups. (F) Proportion of large, medium, and small P-bodies in *vps1*Δ cells across the cell size groups.

This discrepancy can be reconciled by considering the nonlinear relationship between volume and linear distance: a 10% reduction in cytoplasmic volume translates to only about 3.45% shrinkage in radius, equivalent to ∼0.17 µm for a typical yeast cytoplasmic dimensions (∼5 µm). This reduction is minor compared to the ∼2.2 µm nucleation distance inferred from simulations. Thus, small changes in cytoplasmic diameter may be insufficient to affect nucleation probability, yet they may impair granule coalescence, favoring fragmentation in *sch9*Δ cells. In addition, the simulated puncta numbers correspond to a scenario in which the central sphere occupies less than 1% of the whole-cell volume, using a d_ₙ_ of 2.2 µm, which is far below the actual occupation by the organelles (Fig. 3D-F). This suggests that effective intervention by the cellular organization must be much smaller, possibly due to the condensate itself, which diffuses within the cell. These findings experimentally confirm that intracellular crowding affects P-body morphology and support the existence of a spatially constrained fusion threshold that governs the organization of condensates.

### Larger P-bodies exhibit delayed disassembly upon nutrient repletion

Having established that spatial parameters regulate condensate number and size, we next asked whether these morphological features influence functional stability. In classical phase-separation systems, larger droplets are thermodynamically favored and dissolve more slowly due to reduced surface-to-volume ratios and kinetic buffering ^40^. We hypothesized that large P-bodies would persist longer upon glucose reintroduction, providing a functional correlation to their size. We performed time-lapse confocal imaging of glucose-starved cells subjected to nutrient repletion and tracked P-body disassembly across a 10-minute window. As expected, small and medium P-bodies disappeared rapidly, with more than 80% disassembling within four minutes (Fig. 5A–B). In contrast, large P-bodies were highly stable, often persisting until the final time point. This size-dependent disassembly rate suggests that larger condensates have altered material properties, such as increased viscosity or reduced surface curvature, that slow their dissolution. These results suggest that condensate size is not merely a geometric consequence of fusion events but has functional implications for granule stability and cellular memory. Larger P-bodies may act as more robust repositories for mRNA and protein complexes, potentially influencing translational reactivation or RNA decay following stress relief. This observation supports a model in which P-body size encodes distinct biophysical states, governed by nucleation and fusion history, and modulated by spatial constraints during formation.

**Figure 5.**
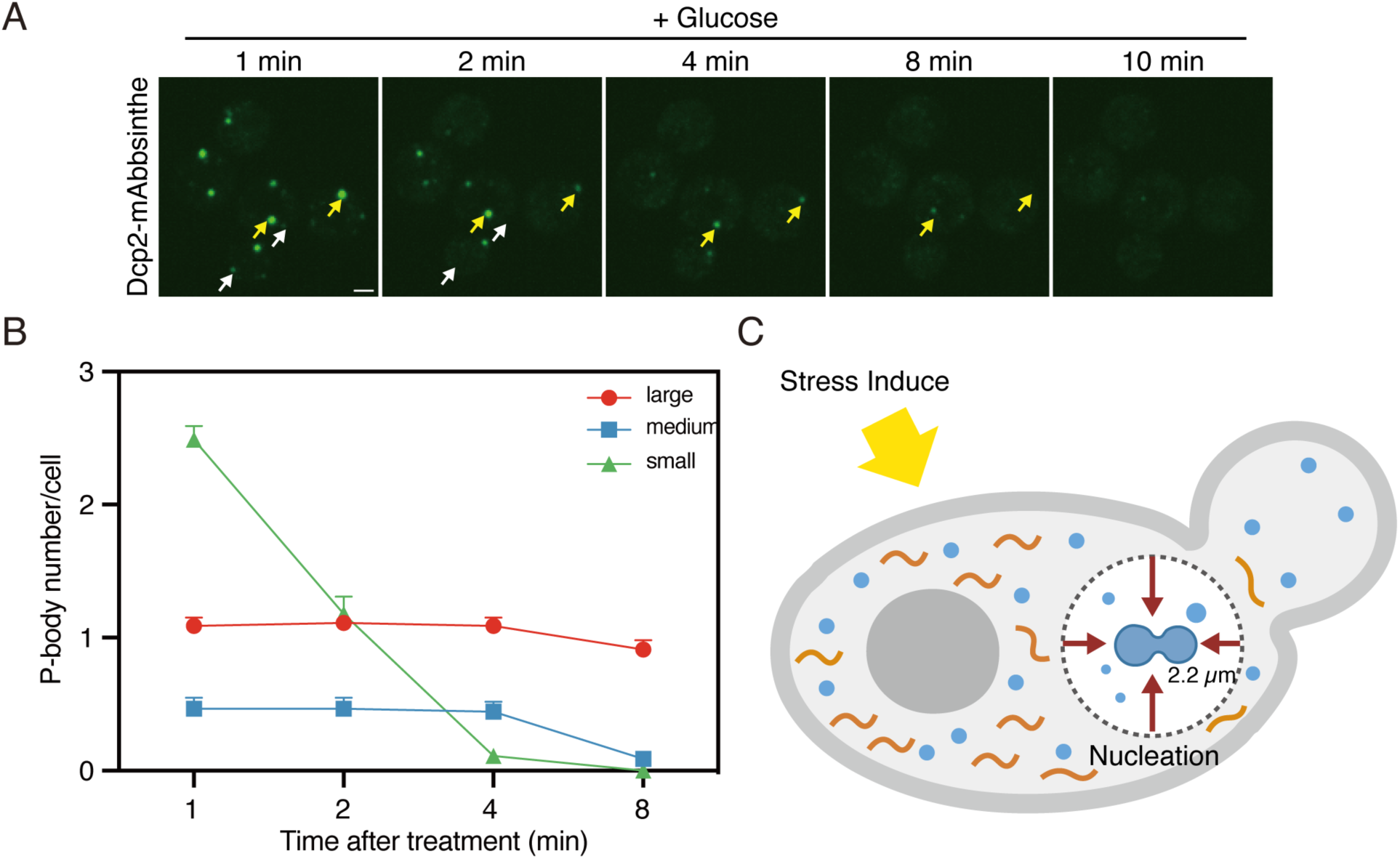
| Larger P-bodies exhibit delayed disassembly upon nutrient repletion (A) Time-lapse images of P-bodies in yeast cells following glucose reintroduction after 30 minutes of starvation. Images were taken at 1, 2, 4, 8, and 10 minutes after the addition of glucose. Yellow arrows indicate large P-bodies, white arrows indicate small P-bodies. Images are z-projections of 42 slices. Scale bar, 2 µm. (B) P-body numbers per cell over time after glucose reintroduction. Small P-bodies rapidly disaggregate, while large P-bodies remain stable. P-body size is primarily determined at the initial time point of formation. n> 45 cells within three independent experiments. Error bar represents s.e.m. (C) Schematic model of P-body formation in yeast cells. Upon stress induction, molecular components (blue dots) and mRNA (orange lines) condense into P-bodies via nucleation within a spatial nucleation distance (∼2.2 µm).

## Discussion

Our quantitative imaging combined with microfluidic cell size sorting and Monte Carlo modeling shows that phase-separated P-bodies follow a characteristic nucleation distance (∼2.2 µm), which determines their abundance as a function of overall cell diameter (Fig. 1, 2). By demonstrating that a single geometric parameter is sufficient to recapitulate the full empirical relationship between cell size and condensate number, we provide, to our knowledge, the first *in vivo* measurement of a nucleation distance for any biomolecular condensate. This finding confirms long-standing predictions from classical nucleation theory ^20–23^ and extends them to the crowded, heterogeneous cytoplasm. Previous work has shown that many organelles scale with growth ^24–29^, but whether equivalent principles govern LLPS compartments has been unclear. Our microfluidic approach allowed us to modulate cell diameter without genetic perturbation or cell cycle arrest, cleanly separating geometric from biochemical variables. The experiments showed larger cells consistently accumulated more, but smaller P-bodies (Fig. 1D). The simulation showed that this pattern naturally arises when nucleation events are prevented within dₙ of an existing droplet: as the cytoplasmic volume increases, the number of non-overlapping spheres of radius dₙ roughly increases with volume (Fig. 2A-C). These data argue that cells can modulate condensate abundance passively, simply by growing. Our minimal model deliberately omits molecular interactions, active transport, or cytoskeletal barriers, yet captures the essential statistics of P-body distribution. This parsimonious view suggests that dₙ signifies a fundamental effective reaction radius that incorporates all microscopic determinants (multivalent interactions, crowding, diffusivity) into a single mesoscale parameter. Future work can refine this framework by allowing dₙ to vary in response to environmental cues, post-translational modifications, or ATP levels, ultimately enabling the quantitative prediction of condensate patterns from first principles.

Extending the model to include a central vacuole and nucleus showed that increasing internal occupancy raises condensate number even at constant cell size (Fig. 3). The effect is conceptually similar to the soft wall confinement used to promote vapour nucleation in microfluidic cavitation chambers: reducing accessible volume raises the local supersaturation of phase separating components and shortens the mean free path required to exceed the nucleation barrier. The *vps1*Δ mutant, which has an enlarged vacuole, validated this prediction in vivo, exhibiting higher P-body counts across all size bins (Fig. 4D). By contrast, *sch9*Δ increased vacuole volume but did not change granule number, highlighting that modest reductions in linear distance (< 0.1 µm) are negligible relative to dₙ, yet still fragment larger condensates and bias the population towards smaller droplets (Fig. 4E–F). Together, these observations position spatial confinement as a tunable knob for condensate biogenesis, complementing known biochemical regulators such as post-translational modifications or RNA scaffolding ^9–12,14–18,41^.

The strikingly slower dissolution of large P-bodies after glucose re-addition (Fig. 5) mirrors classic Ostwald ripening behaviour, in which surface curvature drives exchange with the soluble pool. Our data, therefore, suggest that the history of nucleation and fusion events leaves a lasting physical imprint that regulates the lifetime of stress-induced mRNA repositories. A speculative but attractive model is that large, stable granules favour selective retention of particular mRNAs or protein complexes, endowing cells with a tunable memory of past stress episodes. P-bodies are known to appear in G1 and grow through G2 without changing in number ^33^. We reconcile this observation with our findings by noting that budding yeast double their diameter only late in the cell cycle; before this size threshold is crossed, the cytoplasm remains too small to permit additional nucleation sites. More broadly, any physiological or pathological state that alters cell volume or organelle allotment, such as hypertrophic growth, vacuolar fragmentation in aging, or lysosomal enlargement in neurodegeneration, could reshuffle the cellular condensate landscape. Indeed, impaired clearance of RNA granules is implicated in ALS and related disorders ^6^. We propose that disrupted spatial scaling, rather than intrinsic changes in phase-separating proteins, may contribute to the aberrant accumulation of condensates in these contexts.

We imposed glucose starvation to synchronize P-body formation; whether dₙ is conserved for other stresses or constitutive granules, such as stress granules, remains to be tested. Additionally, the model treats cells as perfect spheres and neglects the geometry of buds or cytosolic viscosity gradients. High-speed light sheet imaging of nascent granules could capture accurate nucleation kinetics and refine these assumptions. Furthermore, our static snapshots cannot distinguish between *de novo* nucleation and the fission of pre-existing droplets; combining selective photoconversion with long-term tracking would resolve this issue. Finally, although we focused on scaffold protein concentration, RNA itself can nucleate P-bodies ^41^; incorporating measured RNA fluxes may sharpen model accuracy. By quantitatively bridging cell-scale geometry with mesoscale phase behaviour, our study uncovers a simple spatial rule that governs how many condensates a cell can host. We anticipate that nucleation distance will prove to be a unifying concept across diverse LLPS assemblies, offering a geometric lens through which to interpret the dynamic self-organization of the cytoplasm.

## Methods

### Yeast strains and plasmids

The yeast strains and plasmids used are listed in Supplementary File 1, and the oligonucleotides used for plasmid construction and gene modification are listed in Supplementary File 2. To reduce variability among the constructed yeast strains, the strains were created either through the integration of a linear PCR product or a plasmid linearized through restriction digestion. The variants of yeast expression plasmids (TTP257, PXQP017) for fluorescent protein were constructed by swapping GFP of pFA6a-link-yoGFP-SpHis5^42^ with mAbb0.5 and mAbbsinthe^38^ through the combination of Gibson assembly and PCR. Fluorescent protein tagging for Dcp2 and Edc3 with mAbb0.5, mAbbsinthe, and yoGFP was performed with PCR-mediated homologous recombination using TTP257, PXQP017, and pFA6a-link-yoGFP-SpHis5, and integrations were confirmed by microscopy. Su9-mCherry was expressed by the TDH3 promoter from the integrated plasmid TTP076, and the strains with three copies of the integrated plasmids in a cell were screened through microscopy and used for further experiments. Deletion mutant strains were constructed with PCR-mediated homologous recombination using pFA6a-hphMX6, and the integration of the hphMX6 cassette was confirmed by PCR followed by electrophoresis.

### Microfluidic device fabrication

The design and fabrication of the microfluidic device for yeast replicative aging were based on previously published work ^43,44^. To enable the capture of yeast cells with different diameters, the initial 6 µm cell trap design was adjusted to include trap diameters ranging from 5.1 to 10 µm.

### Setting up a microfluidic experiment

Before the microfluidics experiment, the device was placed under vacuum for 20 minutes to remove air from the channels. The device was then placed on the stage of an inverted spinning disk confocal microscope. Media ports were connected to plastic tubing, which in turn was connected to 20 mL syringes filled with fresh synthetic complete medium (SD) (CSM powder from Sunrise Science, #1001-100, supplemented with 2% glucose and 0.04% Tween-20). These syringes were positioned approximately 24 inches above the microscope stage to drive media flow via gravity. Yeast cells were grown in SC medium at 30℃ with a rotator speed of 70 rpm until they reached an OD_600_ of approximately 1.0. The following day, yeast cells were diluted to an OD_600_ of 0.1 and cultured for an additional 4 to 5 hours until they reached an OD_600_ of approximately 0.6, before being used in the experiment. To capture the cells in the microfluidic device, the culture was diluted 4-fold and transferred into a 20 mL syringe prefilled with fresh SC medium containing 0.04% Tween-20. The media syringe was replaced with the syringe containing the yeast culture temporarily for loading cells. Medium flow into the device was gravity-driven, allowing cells to enter and occupy the traps. After loading, the yeast culture syringe was replaced with the media supply syringe, and all tubing heights were adjusted to maintain a height differential of approximately 60 inches.

### Microscopy

Cells were imaged at 30℃ by an Eclipse Ti2-E Spinning Disk Confocal with Yokogawa CSU-X1 (Yokogawa) with 50 µm pinholes, located at the Tsinghua-SIGS Tsuboi Laboratory. Imaging was performed using a CFI Plan Apochromat Lambda 100x 1.49 NA oil objective. Z-stacks (300nm steps) were acquired by a Prime 95B sCMOS camera (Photometrics) (pixel size 0.11µm). Imaging was controlled using NIS-Elements software (Nikon). Images of the microfluidic device were acquired using the large-image acquisition mode. The scanning area was set to 0.1 × 1.5 mm with a 10% overlap between adjacent fields.

A P-body visualization experiment in a 96-well plate was performed as follows: Yeast cells were grown in SC medium with 2% glucose at 30°C with a rotator speed of 70 rpm. 100μl of mid-log phase yeast cells (OD600 of 0.4 to 0.7) were harvested and placed into a 96-well Glass Bottom Plate (Cellvis LLC) coated with 0.1 mg/mL concanavalin A (Sigma-Aldrich C2010). After 15 minutes of incubation, the well was washed twice with 100 μL of SC media without glucose and then filled with 100 μL of SC media without glucose.

### The measurement of the photobleach resistance

To measure the photobleach resistance of GFPs, a 488 nm laser and a 510–520 nm emission filters were used. The cells were imaged with 50% of pre-set laser power 20 times. 25 frames of the z-section with a 100-millisecond exposure time were acquired in steps of 300 nm. After background subtraction, the total fluorescence intensity was divided by the cell occupation area in the image. The fluorescence intensity was normalized to the initial intensity to obtain Relative Fluorescence Strength (RFS). The average RFS of the 30 cells was calculated, and the RFS curve was created by MATLAB (version 2021a) using polynomial curve fitting.

### Molecular nucleation model for P-body formation

The model was based on a Monte Carlo simulation approach. Since the yeast cell is approximately spherical, it was modeled as a spherical region in the simulation. P-bodies are randomly distributed in the cytoplasm and are also roughly spherical; therefore, in the simulation, they were also defined as spherical molecules. Details were as follows:

First, the initial number of molecules per cell and the number of simulated cells were defined. Cell diameter was used to represent different cell sizes. In each simulation, the initial position and volume of each molecule were assigned using a random number generator, and their positions were marked in a three-dimensional Cartesian coordinate system. The following procedure was employed to generate the three-dimensional coordinates for each molecule:

1. Random angle generation: Two angles, θ and φ, representing the azimuthal and polar angles in spherical coordinates, were generated using uniform random distributions.
2. Radial distribution: A radial coordinate was calculated based on the predefined cell radius, with a cube root distribution applied to ensure uniform volume distribution.
3. Position conversion: Using the standard spherical-to-Cartesian coordinate transformation, the position in spherical coordinates was converted to three-dimensional Cartesian coordinates according to the following equations:

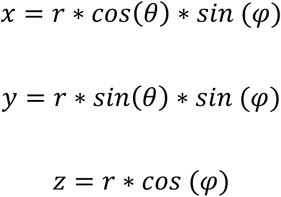

Since this study induced P-body formation by applying glucose starvation stress, the mobility of P-bodies in this condition is very slow and can be neglected ^45^; therefore, the molecules were considered to be in a static state. When the distance between two molecules fell below a critical threshold (a predefined nucleation distance), these molecules were assumed to aggregate to become one molecule.

The aggregation process was implemented as follows:

1. Distance calculation and collision detection: To efficiently identify the closest pair of molecules, an auxiliary function, *findclosestpoint*, was developed. Different models use different *findclosestpoint* functions, as mentioned in Figure 3.
2. Molecule merging: Upon collision, the volumes and positions of the two molecules were merged. The resulting molecule had an increased volume, and its position was determined by *findclosestpoint*.
3. State update: The number and positions of molecules in the cell were updated accordingly after each aggregation event.

The code updated the distribution of molecules in the cell based on the current positions and volumes at each simulation step, progressively approaching the final aggregation state. The simulation loop was terminated once the distances between all molecules exceeded the predefined nucleation distance.

### Quantification and statistical analysis

Detailed statistics, including the number of cells analyzed, mean value, standard deviation, and standard error of the mean, are indicated in each figure legend. The Wilcoxon rank-sum test was performed using GraphPad Prism (GraphPad software).

## Data and materials availability

All data supporting the findings of this study are available within the paper and its supplementary information. Yeast strains and plasmids used in the study are provided in Supplementary File 1. The oligonucleotides used are listed in Supplementary File 2. Further information and requests for raw data and reagents should be directed to and will be fulfilled by the lead contact, T.T.

## Code Availability

The Pytorch codes for p-body drug screening can be found at https://github.com/xuexuegu/nucleation.git.

## Supplemental information

Supplementary File 1 | List of yeast strains and plasmids Supplementary File 2 | List of oligonucleotides

## Acknowledgments

We thank members of the Tsuboi laboratory, especially Abdul Haseeb Khan, for helpful discussions and feedback on the paper. We thank Nathan Shaner and Gerard Lambert for sharing mAbb0.5 and mAbbsinthe plasmids and Nan Hao for sharing the design of the microfluidic device. This work was supported in part by the Key Research and Development Program of the Ministry of Science and Technology 2024YFE0102700, 2023YFA0914303, Shenzhen Science and Technology Innovation Commission WDZC20220811144737001, startup fund OD2021031C, Interdisciplinary Research and Innovation Fund JC2022008, and Overseas Research Cooperation Fund HW2024009 from Tsinghua SIGS (to T.T.),

## Author contributions

X.G. and T.T. conceived and designed the project. X.G. and X.Q. performed wet experiments and analyzed image data. X.G. and D.S. performed image quantification and computational analysis. X.G. and T.T. wrote the manuscript.

---

**Supplementary Figure 1.**
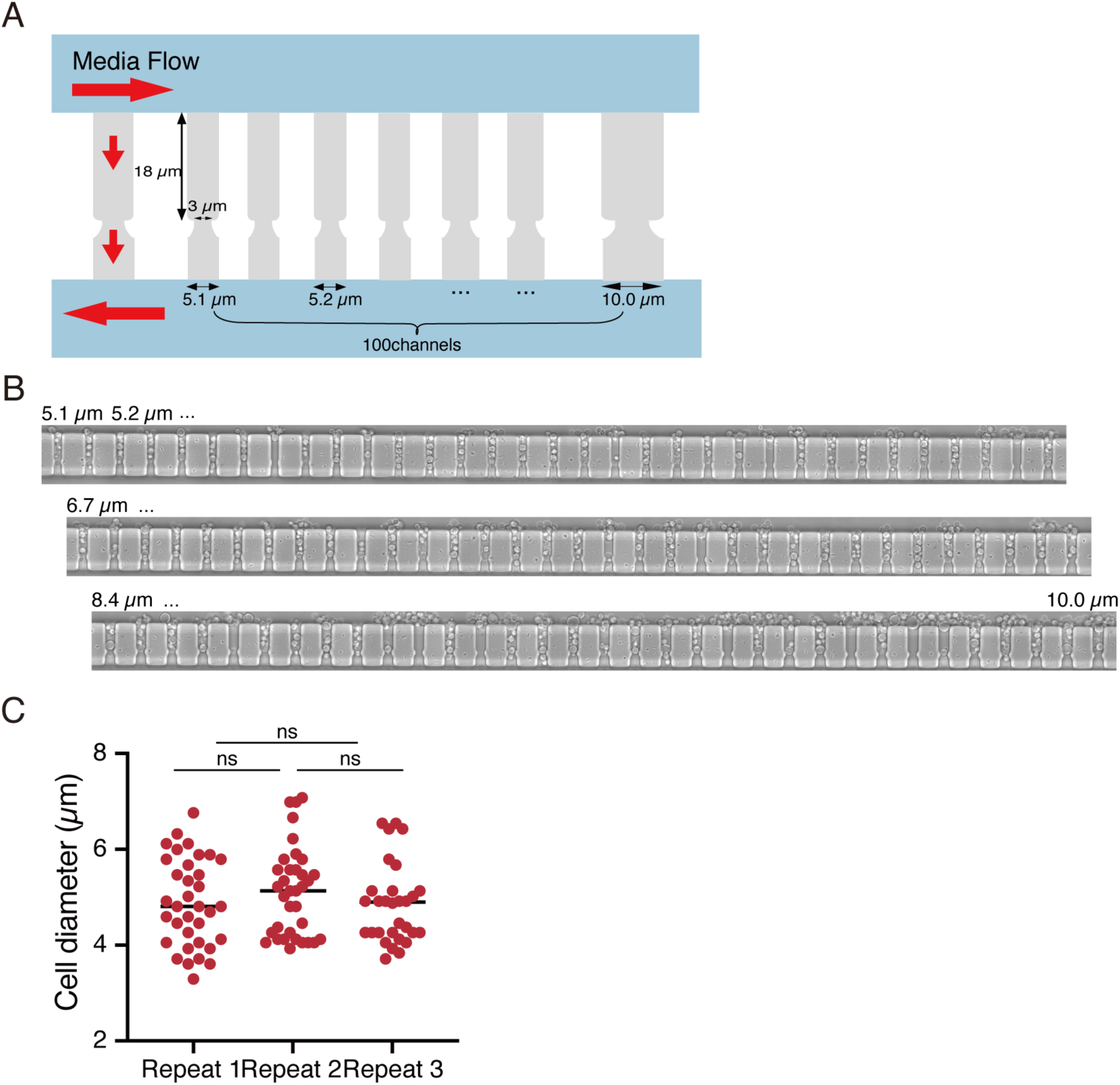
| Design of microfluidics for different sizes of the cells (A) Schematic design of the microfluidics device for capturing yeast cells of different sizes. (B) Representative microfluidic device with a well size spanning 5.1 µm to 10 µm. (C) The microfluidics consistently captures cells of different sizes (n ≥ 30 cells per repeat).

**Supplementary Figure 2.**
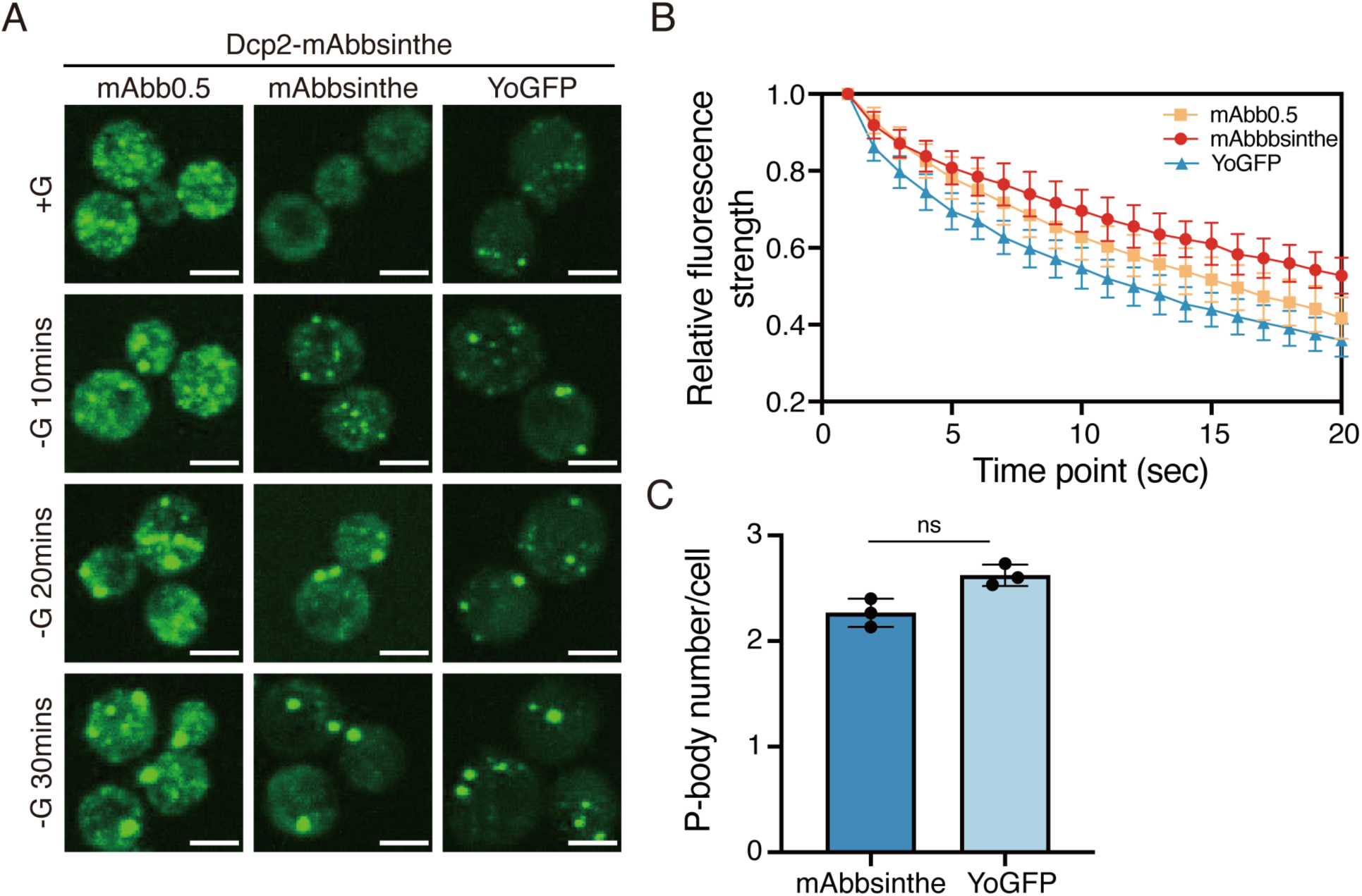
| Dcp2 labeled with mAbbsinthe exhibits similar condensation and greater resistance to photobleaching compared to GFP (A) Z-stack images of Dcp2 labeled with mAbb0.5, mAbbsinthe, and YoGFP. Cells were visualized during glucose (G) starvation at 0, 10, 20, and 30 minutes. All images share identical contrast settings. The scale bar indicates 2 µm. (B) Various Dcp2-GFPs were photobleached under a 10-Watt laser for a 50-ms exposure for each time point. The fluorescence recovery was monitored for 20 seconds (n = 30 cells) and the relative fluorescence strength was plotted. (C) P-body numbers per cell for mAbbsinthe and YoGFP-tagged Dcp2 after glucose starvation for 30 min. Statistical significance was assessed by the Unpaired T-test (ns P > 0.05, n>30).

**Supplementary Figure 3.**
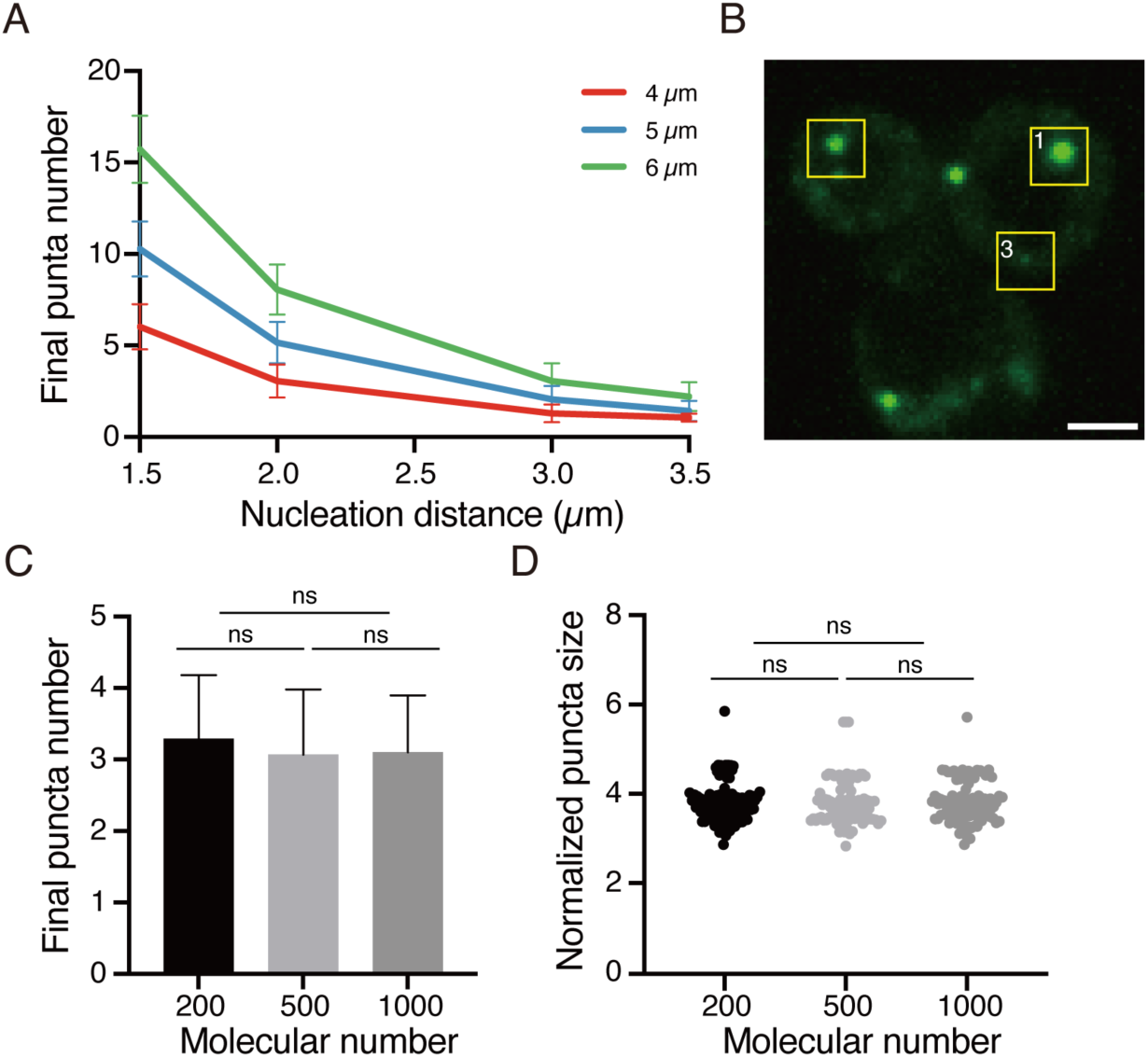
| Effects of nucleation distance and molecular number on P-body formation simulation. (A) Relationship between the final puncta number across different nucleation distances with different sizes of the cell. (B) Representative images of the three P-body size classes. Yellow boxes highlight the representative P-bodies in the different size groups. 1 - large, 2 - medium, and 3 - small groups, respectively. (C) Final puncta number is independent of molecular concentration in the nucleation simulation. The bar graph shows the final number of simulated P-bodies when varying the initial number of molecules (200, 500, 1000). Statistical significance was assessed by the Unpaired T-test (ns P > 0.05, n = 200 simulations per group). (D) Puncta size distribution does not change upon molecular number. Statistical significance was assessed by the Unpaired T-test (ns P > 0.05, n = 200 simulations per group).

