## Supplemental File 1 for "A nucleation distance sets the cell size-dependent P-body assembly"

### Supplementary File 1. Yeast strains and plasmids used in this study

| Strain/plasmids | Genotype[plasmid](plasmid number) | Source |
|---|---|---|
| **Strains** | | |
| W303-1a | MATa *ade2-1 can1-100 his3-11 leu2-3 trp1-1 ura3* | Lab stock |
| XFY082 | W303-1a sch9Δ::hphMX | This study |
| XFY083 | W303-1a vps1Δ::hphMX | This study |
| **Plasmids** | | |
| pNCST-mAbb0.5 | | This study |
| pNCST-mAbbsinthe | | This study |
| pFA6a-link-yoGFP-SpHis5 | | Lee et al., 2013 |
| TTP257 | pFA6a-link-mAbb0.5-SpHis5 | This study |
| PXQP017 | pFA6a-link-mAbbsinthe-SpHis5 | This study |
| TTP076 | pRS406*GPD*p-Su9-mCherry | Tsuboi et al.,2020 |
| ZP191_pFA6a-hphMX6 | | Tsuboi et al.,2020 |
