## Supplemental File 2 for "A nucleation distance sets the cell size-dependent P-body assembly"

### Supplementary File 2. List of oligonucleotides used for plasmid construction

| Gene Name | Oligonucleotides used for plasmid construction |
|---|---|
| *GA_KT_mAbb0.5Q_F* | 5'- CGGTGACGGTGCTGGTTTAATTAACATGGTGAGCAAGGGCGAGGAG -3' |
| *GA_KT_mAbb0.5Q_R* | 5'- TCGCTTATTTAGAAGTGGCGCGCCTCACTTCTGCAGCTCGTCCATGCC -3' |
| *mAbbsinthe_insert_F* | 5'- GTCGACGGATCGGTGACGGTGCTGGTTTAATTAACATGGTGAGCAAGGGCGAGG AGCTCTTC -3' |
| *mAbbsinthe_insert_R* | 5'- CATAAGAAATTCGCTTATTTAGAAGTGGCGCGCCTTACTTGTACAGCTCGTCCAT GCCGTTGATAGC -3' |
| *Dcp2_GFP replacement F* | 5’- TTACAGTGTGTCTATAAAACGTATAACACTTATTCTTTCATCGATGAATTCGAG CTCG -3’ |
| *Dcp2_GFP replacement R* | 5’- CGATGACAAGCCATTGTACAAGAGAATGTTCAGCACCAAAGGTGACGGTGCTGG TTTA -3’ |
| *Edc3_GFP replacement F* | 5’- TGATCTTTTCGTCACTGACGGGTCCCTGCTATTAGATTTGggtgacggtgctggttta-3’ |
| *Edc3_GFP replacement R* | 5’- ACGTATGTATCCAGTTTAGGCTAAAGTAATTCTTGGTTTAtcgatgaattcgagctcg-3’ |
| *Vps1D::MX4/6-F* | 5'- ATAAGGACCGTACGAAAACTGCACATTTTATATTATCAGATATC cggatccccgggttaattaaggcg-3' |
| *Vps1D::MX4/6-R* | 5'-GAAATACTCAAAACCAAGCTTGAGTCGACCGGTATAGATGAGGAAAC cataggccactagtggatctga-3' |
| *Sch9 D::MX4/6-F* | 5'- AGAATTATACTCGTATAAGCAAGAAATAAAGATACGAATATACAAT cggatccccgggttaattaaggcg-3' |
| *Sch9 D::MX4/6-R* | 5'- AAAGAAAAGGAAAAGAAGAGGAAGGGCAAGAGGAGCGATTGAGAAA cataggccactagtggatctga-3' |
